# MetAlyzer to Perform Streamlined, Interactive, and Pathway-Mapping Analysis of Targeted Metabolomics Data from the biocrates Platform

**DOI:** 10.64898/2025.12.17.692011

**Authors:** Qian-Wu Liao, Luis Herfurth, Christina Schmidt, Nils Mechtel, Hagen M. Gegner, Alice Limonciel, Marc A. Schneider, Julio Saez-Rodriguez, Rüdiger Hell, Junyan Lu, Gernot Poschet

## Abstract

Mass spectrometry (MS)-based metabolomics has emerged as a powerful tool to address multifaceted biological questions. Commercial solutions like the ones developed at biocrates allow reliable and quantitative targeted metabolic profiling, including the conversion of the raw MS spectra into absolute concentrations of metabolites. These results can be exported for further analysis under several formats with varying levels of human-vs. machine-readability. The default output format is an Excel spreadsheet that favours human readability and therefore requires extra preparation steps for downstream bioinformatic analysis and data exploration. To streamline this next step for users of this platform, we developed MetAlyzer, an R package (https://github.com/Lu-Group-UKHD/MetAlyzer) specifically designed to handle the spreadsheets generated by WebIDQ, the biocrates workflow manager software. MetAlyzer converts WebIDQ-generated spreadsheets into flexible *SummarizedExperiment* objects and provides functions for data preprocessing, statistical testing, and visualization of differential metabolites. To further support data exploration and hypothesis generation by users without coding experience, we also developed an interactive and intuitive Shiny app (https://metalyzer.shinyapps.io/MetAlyzer_ShinyApp/) that interfaces with MetAlyzer’s core functionality, enabling users to execute the complete analysis workflow without writing code. This combination can help scientists deepen their understanding of metabolomics results, supporting the broader adoption of metabolomics in the life sciences community.

## Introduction

Metabolomics, aided by mass spectrometry (MS) technologies, has become a powerful tool for investigating intricate biochemical networks within living organisms and is applied across diverse fields, such as biomedical, nutritional, and agricultural sciences (Gowda et al., 2008; Gomez-Casati et al., 2013; Gonzalez-Covarrubias et al., 2022). While global metabolic profiling has attracted significant interest over the past decades due to its comprehensive coverage and discovery potential, targeted metabolomics offers unique advantages in specificity, sensitivity, and reproducibility by focusing on the precise measurement and quantification of predefined sets of metabolites (Begou et al., 2017).

To achieve reliable MS-based metabolomics analysis, biocrates, a biotechnology company, has designed ready-to-use kits that provide quantitative, standardized, and reproducible assays targeting several hundred or more than 1,000 metabolites across diverse metabolite classes, depending on the respective kit version (biocrates, Austria). These kits have been used in studies involving a wide range of organisms and sample types (Erben et al., 2021; Gegner et al., 2022). Raw MS data from these kits are typically quantified and preprocessed by biocrates’ WebIDQ, a proprietary workflow manager software that transforms MS raw intensities into precise metabolite concentrations. Results can be exported under several formats with varying levels of human-vs. machine-readability. The default output format is an informative but complex Excel spreadsheet favouring human readability and requiring tailored data preparation for downstream statistical analysis and interpretability. Therefore, we created a user-friendly tool that can convert this information-rich but unstructured data into a common data structure (e.g., a data frame), enabling easy integration of these large biocrates datasets for bioinformaticians and wet-lab biologists.

Several web-based platforms can process and analyze targeted metabolomics data, such as MetaboAnalyst (Pang et al., 2024), Galaxy (The Galaxy Community, 2024), and Metabolomics Workbench. However, none of these tools can be directly applied to spreadsheets exported from WebIDQ. MeTaQuaC (Kuhring et al., 2020), a published R package, was developed to perform automated quality control of targeted metabolomics data, particularly from the biocrates platform, and produces reports in HTML format. However, MeTaQuaC and its static reports lack the flexibility for customized data processing and analysis required to answer specific biological questions.

To overcome these limitations, we developed **MetAlyzer**, which aims to facilitate the parsing of WebIDQ outputs, enable flexible analysis, and improve the accessibility of targeted metabolomics data from the biocrates platform. With a dual-mode design that combines a code-based R interface and an interactive Shiny interface (https://metalyzer.shinyapps.io/MetAlyzer_ShinyApp/) within a single package, MetAlyzer accommodates both experienced bioinformaticians as well as wet-lab biologists and medical researchers with limited coding experience. In conjunction with the biocrates platform, we envision MetAlyzer as a complementary and powerful tool for advancing accessible and reproducible metabolomics research in the life sciences. We demonstrate the usability and functionality of MetAlyzer using two datasets: (1) a demo dataset generated with the MxP® Quant 500 XL kit, provided by our collaborators at biocrates, and (2) a published lung adenocarcinoma dataset produced with the MxP® Quant 500 kit (Gegner et al., 2024), as presented in the Results section.

## Results

### Overall features

MetAlyzer provides a workflow for processing, quality control, statistical analysis, and visualization of targeted metabolomics datasets generated using the biocrates platform (Fig. 1). The workflow begins by converting an Excel spreadsheet (.xlsx format) exported from the WebIDQ software into a *SummarizedExperiment* object (Morgan et al., 2022), which serves as the central data structure for subsequent preprocessing and downstream analysis. MetAlyzer supports sample and metabolite filtering, half-minimum imputation, log_2_ transformation, and various analyses including descriptive statistics, fold changes (FCs), and t-tests. Visual summaries of differential metabolites are generated as volcano plots, scatter plots, and metabolic network diagrams. In addition to the R-based interface, MetAlyzer includes a Shiny-based graphical interface (Chang et al., 2024) that offers point-and-click access to its core functionality. The app delivers real-time visual feedback for each preprocessing operation, including boxplots of metabolite concentration distributions across samples and barplots of metabolite missing values and quantification statuses. All preprocessing parameters and their corresponding outputs are logged in a history table, enabling traceability and comparisons of different configurations. For statistical analysis, volcano plots allow users to adjust log_2_ FC and q-value cutoffs and highlight specific metabolites or metabolic classes, while scatter plots show associations between log_2_ FCs and metabolic classes, making it easier to spot differential metabolites and metabolic classes.

**Fig. 1:**
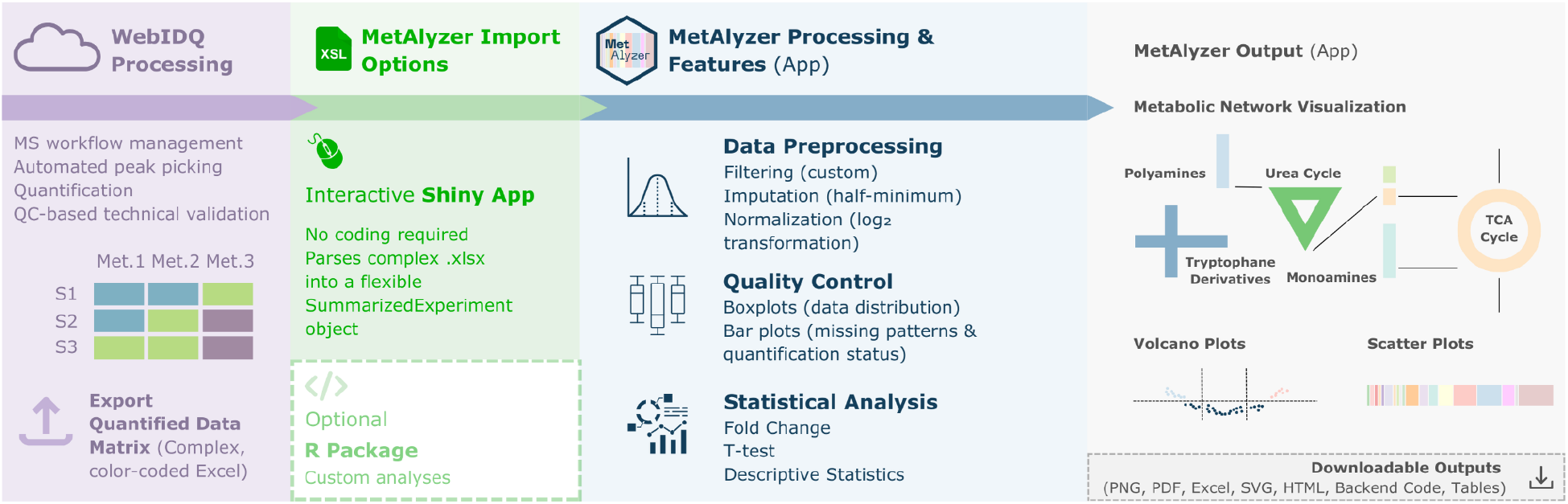
Workflow and overall features of MetAlyzer. The workflow of MetAlyzer begins by converting a WebIDQ-exported spreadsheet into a *SummarizedExperiment* object to which sample and metabolite filtering, data imputation and transformation, and various analyses, including descriptive statistics, fold changes, and t-tests are applied. Although several processing steps in MetAlyzer and WebIDQ appear similar, they serve distinct purposes. For example, WebIDQ normalization focuses on enhancing data accuracy by reducing variability at the sample, batch, and laboratory levels, whereas MetAlyzer normalization aims to improve data normality for downstream statistical analyses. The statistical results can be visualized using volcano and scatter plots and network diagrams. Users can access the full analysis workflow through the Shiny app for point-and-click convenience or customize their analyses using the code-based R version of MetAlyzer.

Another standout feature of MetAlyzer is its contextual visualization of metabolite-level statistics through network diagrams. These diagrams map log_2_ FCs and p-values directly onto curated canonical metabolic pathways that cover the biocrates kits content, such as the TCA cycle, β-oxidation, and glutamate metabolism, which facilitates the identification of biologically relevant metabolites and their interconnections. When a pathway metabolite is represented by multiple derivatives, their statistics are averaged (Supplementary Methods). All visualizations supported by the app are interactive and can be exported in HTML, PDF, SVG, or PNG formats.

### Usage scenario

We demonstrate the utility of MetAlyzer through its Shiny app using a demo dataset generated with the MxP® Quant 500 XL kit, provided by our collaborators at biocrates. The dataset consists of two sample groups, Group 1 and Group 2. The raw dataset exported directly from the WebIDQ software was uploaded into the app and initially visualized through its distribution, missingness patterns, and quantification status composition to assess quality and guide the selection of appropriate preprocessing steps. Given that the overall quantification appeared stable and no systematic distributional drift was evident (Fig. 2A,B), we applied metabolite filtering based on the 80% rule (Wei et al., 2018) and half-minimum imputation to enhance statistical robustness, followed by log_2_ transformation to stabilize variance across the dataset (Supplementary Fig. 1). We then performed statistical analysis comparing the samples from Group 1 with Group 2, which identified differentially abundant metabolites, particularly within the triacylglycerol (TAG) and glycerophospholipid (GPL) classes (Fig. 2C,D). The volcano and scatter plots indicated higher enrichment of TAGs in Group 2, while GPLs were more enriched in Group 1. Finally, the network diagram highlighted potential perturbations in β-oxidation, polyamine, and indole/tryptophan metabolism (Fig. 2E), placing individual metabolites into their functional pathway context (Supplementary Fig. 2; Supplementary Tbl. 1). To demonstrate MetAlyzer’s ability to capture underlying biological signals, we additionally analyzed a published lung adenocarcinoma dataset generated with the MxP® Quant 500 kit (Gegner et al., 2024), with results shown in Supplementary Fig. 3.

**Fig. 2:**
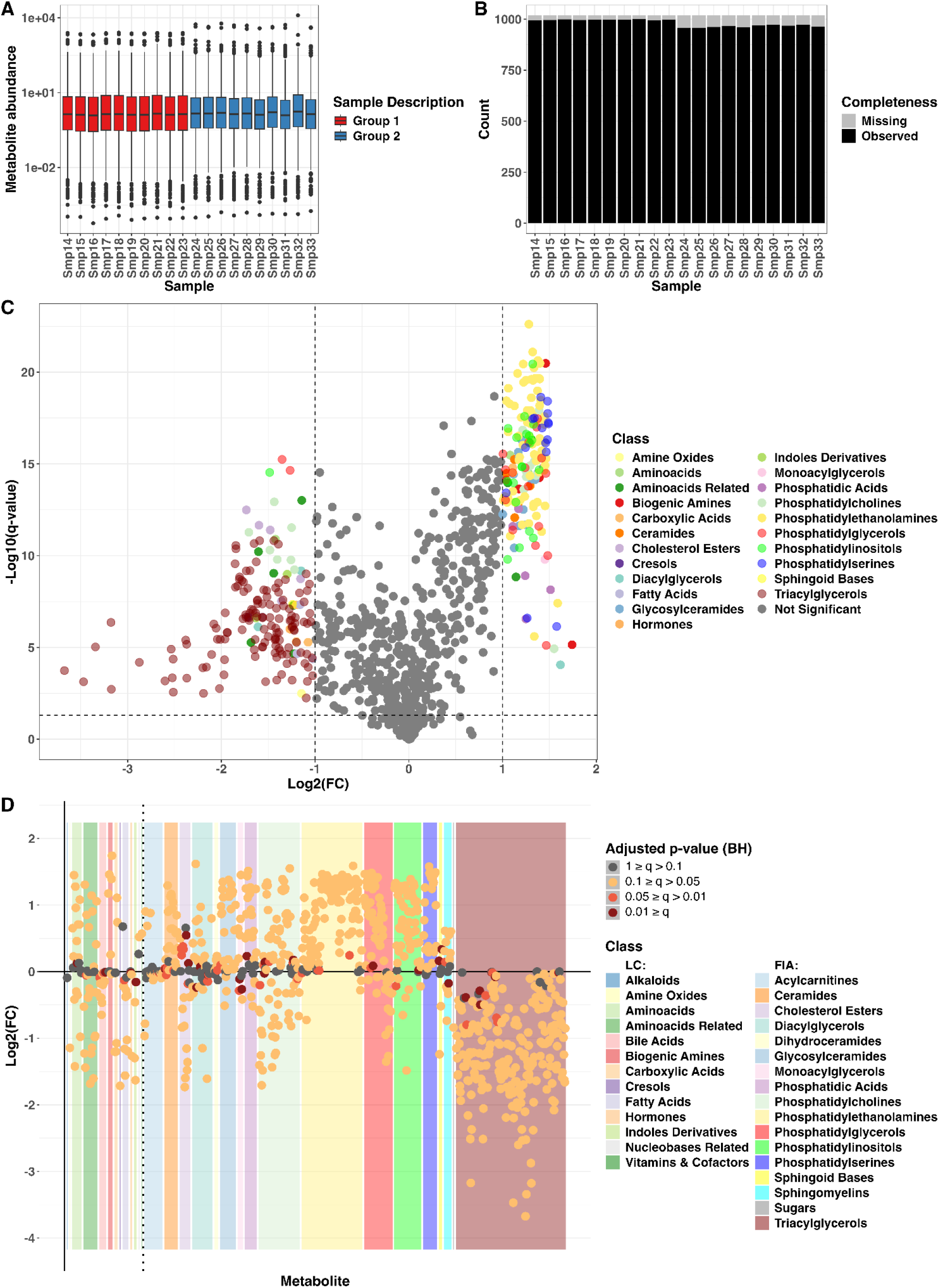
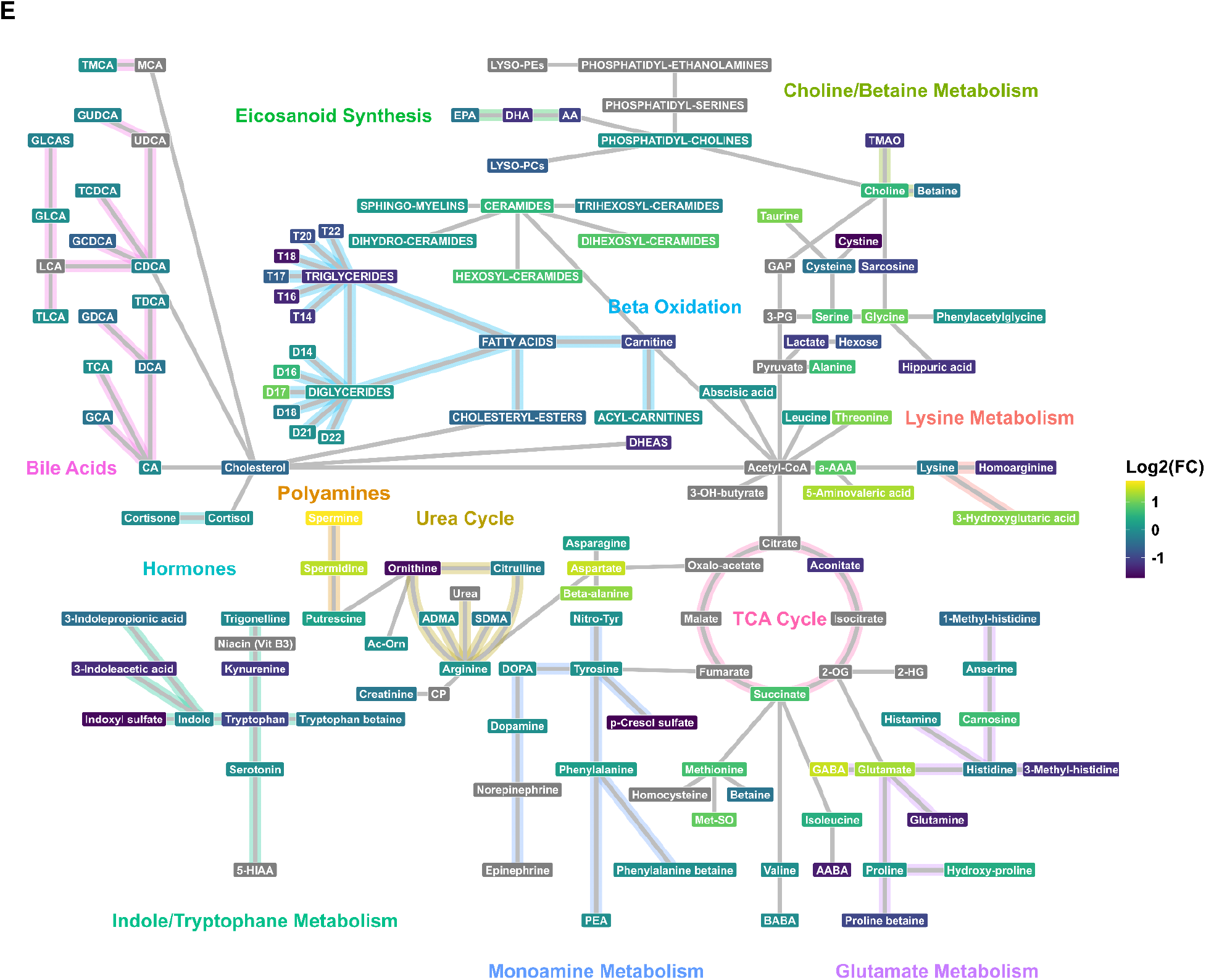
Utility of MetAlyzer demonstrated using demo dataset consisting of two biologically different sample groups. **(A)** The boxplot of the unprocessed data, colored by the two sample groups, showed substantial variance across the dataset but no evident systematic distributional drift. The central line of each box is the median; the box hinges span the 25^th^ to 75^th^ percentiles, while the whiskers extend up to 1.5 times the interquartile range beyond the hinges. Outliers are displayed as individual points outside the whiskers. **(B)** The barplot illustrated missingness patterns across samples in the unprocessed data, suggesting that the overall quantification was stable. **(C)** The volcano plot identified differentially abundant metabolites in the comparison of Group 1 and Group 2 samples, particularly within the triacylglycerol (TAG; brown) and phosphatidylethanolamine (PE; yellow) classes. TAGs were more enriched in Group 2 and PEs in Group 1. The thresholds for log_2_ fold change (FC) and q-value were set at 1 and 0.05, respectively. Colors assigned to points denote metabolic classes. **(D)** The scatter plot complements the volcano plot, illustrating differentially abundant TAGs and GPLs (e.g., PAs, PCs, PEs). Points are colored by q-values; background rectangles indicate metabolic classes. Users can easily view metabolite statistics and class information by hovering over points and background regions in the Shiny app. **(E)** The network diagram revealed potential disruptions in β-oxidation, polyamine, and indole/tryptophan metabolic pathways. Colors of nodes and edges denote log_2_ FCs of metabolites and indicate their involvement in specific pathways.

## Discussion

MetAlyzer is an R package designed for the efficient processing and analysis of targeted metabolomics datasets exported from the biocrates platform. Its strength lies in a streamlined, dual-mode workflow that integrates both a code-based interface and an interactive Shiny app. For users concerned with data privacy, the Shiny app can be readily initialized locally with a single R function call, thereby avoiding reliance on a server-hosted deployment. The workflow begins by converting a WebIDQ-exported Excel spreadsheet containing multiple tabs and colour-coding into a flexible *SummarizedExperiment* object (Morgan et al., 2022), which serves as the central data structure within MetAlyzer and can also be widely used in other external environments (e.g., Bioconductor; Huber et al., 2015), enhancing interoperability and customizability. The workflow is minimal yet functionally complete, encompassing sample and metabolite filtering, data imputation and transformation, statistical analysis, and result visualization, which supports rapid quality control and downstream analysis. Although WebIDQ has its own data cleaning method (modified 80% rule; Wei et al., 2018) and imputation strategies based on the type of missingness (e.g., missing at random or not at random), MetAlyzer provides complementary options, such as filtering by metabolite quantification statuses and imputation using the half-minimum approach.

To improve accessibility and efficiency, the Shiny-based interface enables full workflow execution without requiring users to code, particularly valuable for wet-lab or medical researchers with limited to no programming experience. Unlike existing web-based tools (Pang et al., 2024; The Galaxy Community, 2024), MetAlyzer directly accepts WebIDQ-exported spreadsheets and includes only essential steps, which reduces complexity while supporting flexible data exploration and hypothesis generation. Recognizing that users may require more advanced analyses, we developed MetAlyzer to be readily connectable to MetaProViz, another code-based R-package for further functional analysis to combine the biocrates feature space with metabolite-specific prior knowledge for enrichment analysis, analyze mapping ambiguities and perform biologically informed clustering (Schmidt et al., 2025). We plan to extend MetAlyzer’s functionality by exporting compatible data formats (e.g., concentration matrices) for use in other web-based tools (e.g., MetaboAnalyst for enrichment analysis; Xia and Wishart, 2010). In addition, as newer biocrates kits such as MxP® Quant 1000 cover a broader range of metabolic pathways and compound classes than those available at the start of MetAlyzer’s development, we plan to extend the scope of the current pathway visualization, which remains primarily tailored to MxP® Quant 500 data and may therefore exhibit a degree of coverage bias.

Altogether, MetAlyzer offers an accessible and efficient solution for the reproducible analysis of targeted metabolomics data generated with the biocrates platform, which balances usability and flexibility through a dual interface that supports both routine tasks and exploratory research across varying levels of user expertise.

## Data and Code Availability

The demo dataset used in this study is publicly available in the GitHub repository *Lu-Group-UKHD/MetAlyzer* at https://github.com/Lu-Group-UKHD/MetAlyzer. The MetAlyzer package and the code used to perform the analysis and generate results in this study are available in the GitHub repository *Lu-Group-UKHD/MetAlyzer* and released under the GPL-3.0 license. The MetAlyzer Shiny app can be accessed via https://metalyzer.shinyapps.io/MetAlyzer_ShinyApp/.

## Additional Information

Vignettes for the code-based and Shiny-based versions of MetAlyzer are available in the GitHub repository *Lu-Group-UKHD/MetAlyzer*. A video tutorial illustrating the Shiny app’s functionality is available at https://youtu.be/Gelrvrt1xQ8.

## Acknowledgement

This study is supported by the BMBFTR (Bundesministerium für Forschung, Technologie und Raumfahrt) funded SMART-CARE project (funding code: 16LW0235, 16LW0233K and 161L0213). The Metabolomics Core Technology Platform University is partially funded by the CellNetworks Core Technology Platform (CCTP) of Heidelberg University. The CCTP is funded in part by the BMBFTR and the Ministry of Science Baden-Württemberg within the framework of the Excellence Strategy of the Federal and State Governments of Germany.

