## Supplementary Information for "MetAlyzer to Perform Streamlined, Interactive, and Pathway-Mapping Analysis of Targeted Metabolomics Data from the biocrates Platform"

### Methods

#### Network node statistics

Nodes in the network diagrams represent metabolites or metabolic classes, and their colors indicate statistics such as $\log_2$ fold change (FC). When a pathway member (node) is linked to multiple derivative metabolites, their statistics are summarized according to the following hierarchy to capture the most informative signal:

1. Mean of significant - If one or more derivatives mapping to the node are statistically significant (e.g., based on q-values), the node value is the mean $\log_2$ FC of those significant derivatives only.
2. Mean of measured - If no derivatives are significant but measurements exist, the node value is the mean $\log_2$ FC of all measured derivatives.
3. NA - If no derivatives were measured or included in the dataset, the node value is recorded as NA.

### Figures

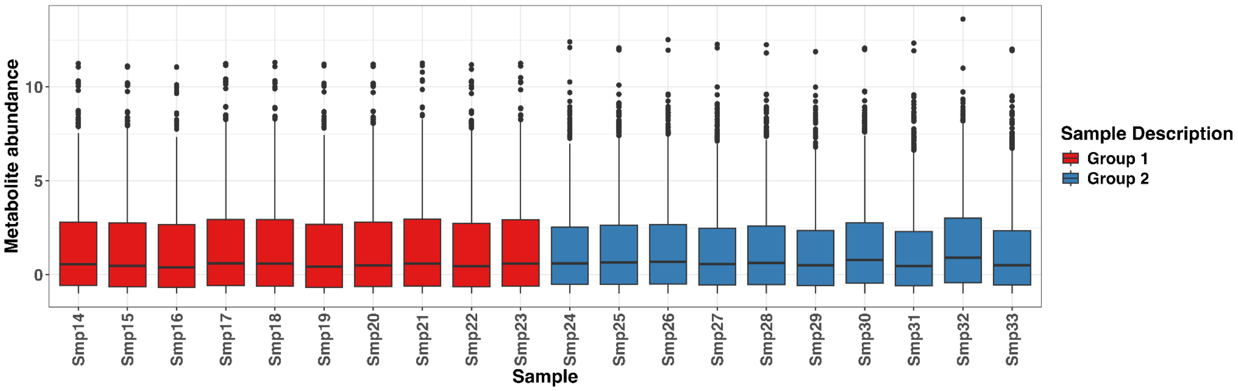


**Supplementary Fig. 1: Distribution of processed demo dataset** | The demo dataset was filtered using the 80% rule, imputed with the half-minimum method, and $\log_2$-transformed to stabilize variance.

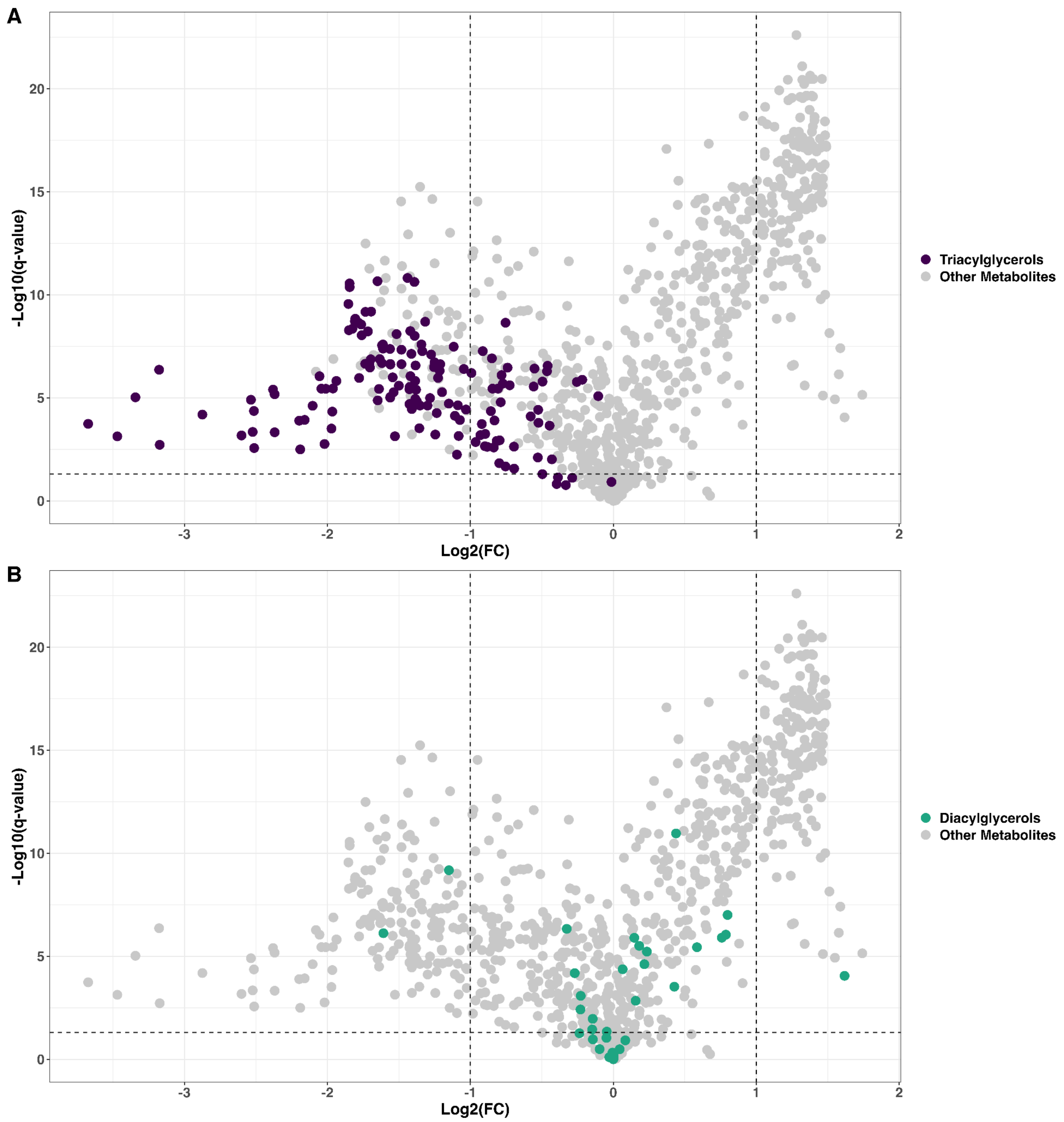


**Supplementary Fig. 2: Volcano plots highlighting Triacylglycerols (TAGs; top) and Diacylglycerols (DAGs; bottom)** | These plots support the pathway patterns shown in the network diagram (Fig. 2E), with TAGs exhibiting negative $\log_2$ fold changes (FCs) and DAGs showing no significant group differences. **(A)** The 151 lipids associated with the T14, T16, and T18 nodes displayed markedly negative $\log_2$ FCs. **(B)** Most of the 40 lipids comprising the D14, D16, and D18 nodes remained largely unchanged between the groups. The thresholds for $\log_2$ FC and q-value were set at 1 and 0.05, respectively.

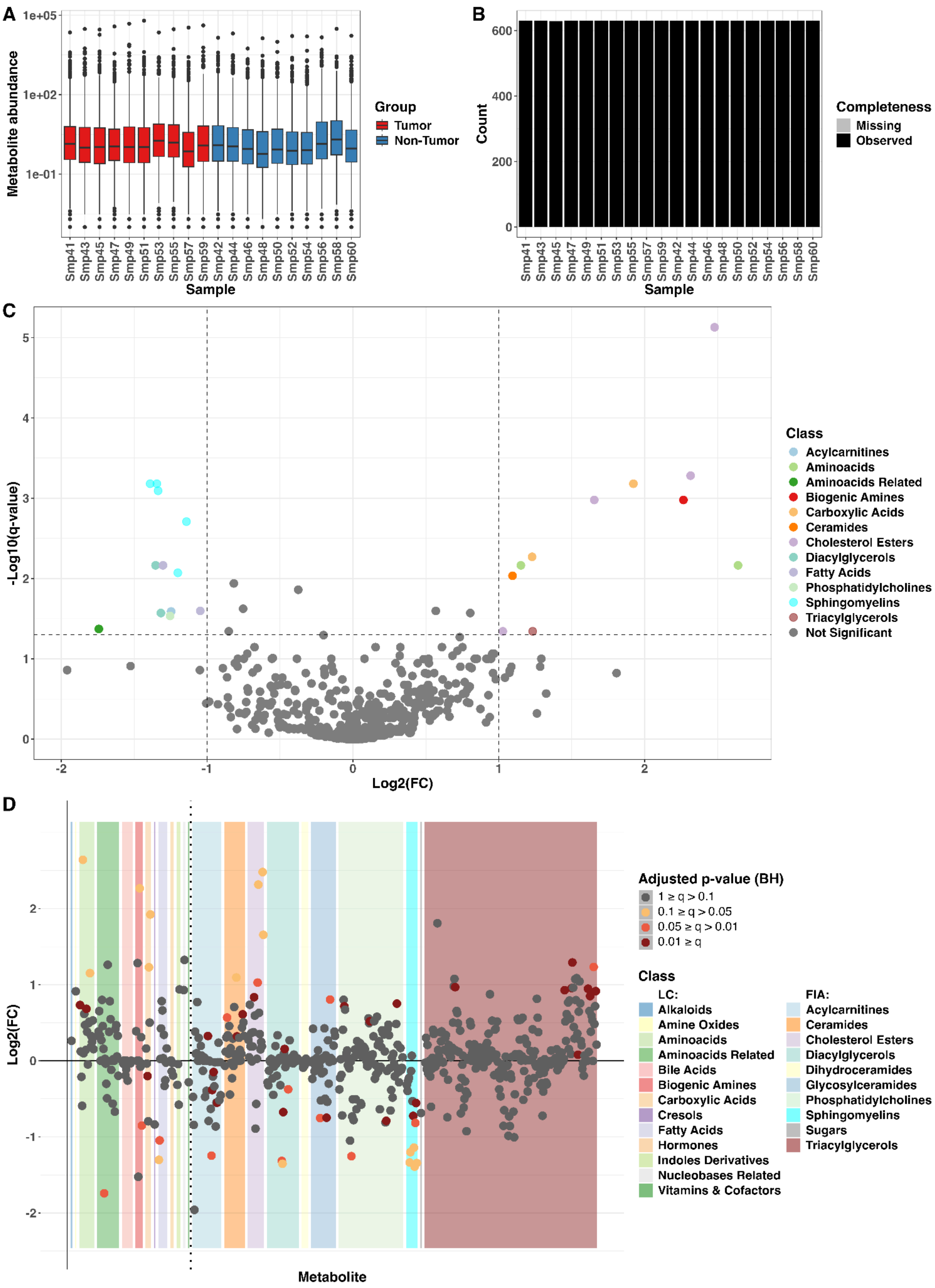
A
Metabolite abundance
1e+05
1e+02
1e-01
Group
Tumor
Non-Tumor
Sample
Smp41 Smp43 Smp45 Smp47 Smp49 Smp51 Smp53 Smp55 Smp57 Smp59 Smp42 Smp44 Smp46 Smp48 Smp50 Smp52 Smp54 Smp56 Smp58 Smp60
B
Count
600
400
200
0
Completeness
Missing
Observed
Sample
Smp41 Smp43 Smp45 Smp47 Smp49 Smp51 Smp53 Smp55 Smp57 Smp59 Smp42 Smp44 Smp46 Smp48 Smp50 Smp52 Smp54 Smp56 Smp58 Smp60
C
-Log10(q-value)
Log2(FC)
Class
Acylcarnitines
Aminoacids
Aminoacids Related
Biogenic Amines
Carboxylic Acids
Ceramides
Cholesterol Esters
Diacylglycerols
Fatty Acids
Phosphatidylcholines
Sphingomyelins
Triacylglycerols
Not Significant
D
Log2(FC)
Metabolite
Adjusted p-value (BH)
1 ≥ q > 0.1
0.1 ≥ q > 0.05
0.05 ≥ q > 0.01
0.01 ≥ q
Class
LC:
Alkaloids
Amine Oxides
Aminoacids
Aminoacids Related
Bile Acids
Biogenic Amines
Carboxylic Acids
Cresols
Fatty Acids
Hormones
Indoles Derivatives
Nucleobases Related
Vitamins & Cofactors
FIA:
Acylcarnitines
Ceramides
Cholesterol Esters
Diacylglycerols
Dihydroceramides
Glycosylceramides
Phosphatidylcholines
Sphingomyelins
Sugars
Triacylglycerols

**E**

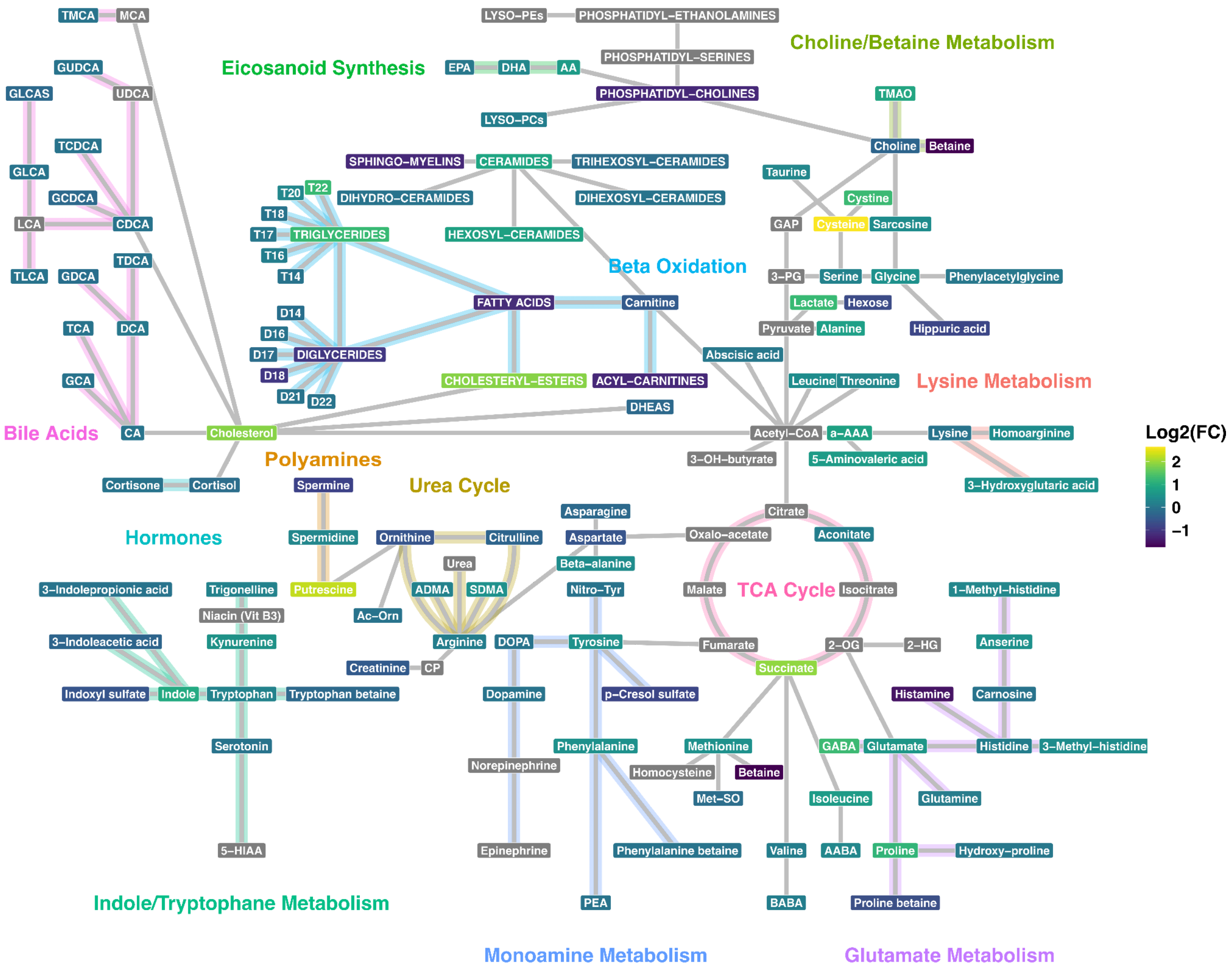


**Supplementary Fig. 3: MetAlyzer analysis of lung adenocarcinoma dataset from Gegner et al., comprising tumor and matched adjacent non-tumor tissue samples** | **(A,B)** Visualization of the raw data using a boxplot and a barplot indicated generally good data quality, with high variance, no apparent systematic drift, and very few missing values (two data points). Consequently, half-minimum imputation followed by $\log_2$ transformation was applied. For the boxplot, the central line is the median; the hinges span the $25^{th}$ to $75^{th}$ percentiles, while the whiskers extend up to 1.5 times the interquartile range beyond the hinges. Outliers are displayed as individual points outside the whiskers. **(C,D)** The volcano and scatter plots highlighted cholesterol esters, cysteine, putrescine, and succinate as significantly enriched in tumor tissue, whereas sphingomyelins, diacylglycerols, and free fatty acids were enriched in non-tumor tissue. These findings are consistent with previous studies on metabolic reprogramming in cancer cells (Xia et al., 2023; Bonifácio et al, 2021; Nowotarski et al., 2013; Eijkelenkamp et al., 2020; Li et al., 2022; Cooke and Kazanietz, 2022). In the volcano plot, thresholds for $\log_2$ fold change (FC) and q-value were set to 1 and 0.05, respectively, and point colors denote metabolic classes. In the scatter plot, points are colored by q-values, and background rectangles indicate metabolic classes. With the Shiny app, users can easily view metabolite statistics and class information by hovering over points and background regions. **(E)** The network diagram identified dysregulation of β-oxidation (Ma et al., 2025; Rashed et al., 2025) and cholesterol esterification, as well as potential perturbations in polyamine metabolism and the TCA cycle, in line with previously reported cancer-associated metabolic reprogramming. Node and edge colors denote $log_2$ FCs of metabolites and their involvement in specific pathways. Collectively, the results demonstrate that MetAlyzer effectively captures underlying biological structure and highlights lipid metabolism-related adaptations in lung adenocarcinoma.

### Tables

**Supplementary Tbl. 1: Statistics of lipid derivatives mapping to T14 node in network diagram** | This table illustrates how $log_2$ fold changes (FCs) for a node linked to multiple metabolite derivatives were summarized. At a significance threshold of 0.05, the $log_2$ FCs of the significant lipids (excluding TG 14:0_39:3) were averaged, yielding a mean of -1.163.

| T14 | | |
|---|---|---|
| **Metabolite** | **Log2(FC)** | **q-value** |
| TG 14:0_32:2 | -1.151 | 1.911e-05 |
| TG 14:0_34:0 | -0.964 | 1.401e-03 |
| TG 14:0_34:1 | -1.300 | 2.380e-05 |
| TG 14:0_34:2 | -1.560 | 2.418e-07 |
| TG 14:0_34:3 | -1.544 | 1.016e-06 |
| TG 14:0_35:1 | -0.430 | 9.665e-03 |
| TG 14:0_35:2 | -0.524 | 1.626e-04 |
| TG 14:0_36:1 | -1.082 | 7.100e-04 |
| TG 14:0_36:2 | -1.717 | 5.956e-09 |
| TG 14:0_36:3 | -1.808 | 1.907e-09 |
| TG 14:0_36:4 | -1.938 | 1.518e-06 |
| TG 14:0_38:4 | -0.526 | 3.823e-05 |
| TG 14:0_38:5 | -0.578 | 7.915e-05 |

| T14 | | |
|---|---|---|
| TG 14:0_39:3 | -0.013 | 1.210e-01 |
